## Supplementary material for "Rift Valley fever virus dynamics in a transhumant cattle system in The Gambia": https://drive.google.com/file/d/1cHUEKmO4qk95uY0lUI6SHzCNIyO6XIvz/view?usp=sharing

Supplementary Text

##### Study Area

This study was conducted across the six administrative regions of The Gambia, an approximately 11,000 km^2^ semi-enclave of Senegal. As the smallest country in mainland Africa, its dominant geographical feature is the Gambia river, which flows for 450km across the elongated country supporting a rich ecosystem including mangrove swamps and wetlands for livestock grazing.

The Gambia has a tropical semi-arid climate with distinct wet and dry seasons. Due to its location at the edge of the Sahara Desert, rainfall is irregular, occurring in a short unimodal season from July to October, with annual rainfall varying between 650 and 1200mm (Touray, 2022). Wet seasons are occasionally characterised by usually heavy rainfall and high humidity. Conversely, the dry season, from November to June, is hot and dry with average temperatures of around 35°C.

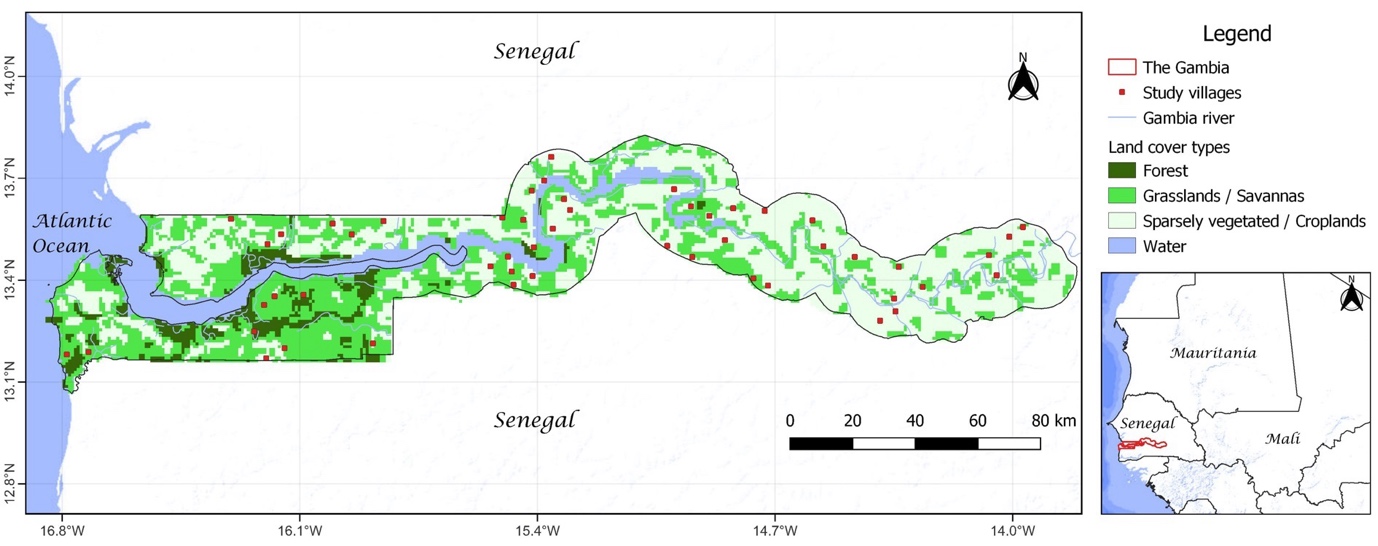
*Figure S1: Map of The Gambia illustrating the location of villages selected for household surveys and ruminant sampling. Map created in QGIS 3.28.3; land cover data derived from Global Land Cover 2000 (GLC2000) Project.*

##### Data Collection

A cross-sectional survey of ruminant livestock linked with household-based questionnaire data was used to gather data on livestock management practices associated with increased odds of Rift Valley fever (RVF) seropositivity across The Gambia (Jarra et al., 2025). Livestock management follows a traditional mixed farming system combining crop husbandry and livestock rearing. Cattle are grazed unrestrictedly on communal resource areas in the vicinity of villages daily, while small ruminants (goats and sheep) are kept near the household compounds. At night, cattle are tethered on the village outskirts, and small ruminants are housed within compounds.

A total of 52 villages were selected using a multi-stage cluster sampling approach from the list of villages in the 2016 National Livestock Census. Within each selected village, up to four livestock-owning households, serving as the primary units of analysis, were randomly selected for participation using the Livestock Census records. Following informed consent, structured interviews were conducted with household heads to gather detailed information about the seasonal movements of their livestock. A pre-tested, standardized questionnaire was used to ensure consistency across study sites, capturing data on seasonal movement (destinations, durations, and routes) of the household animals during the dry season (November-June). To facilitate effective communication and ensure that participants could express themselves comfortably, discussions were held in the local language of the participant.

##### Network Analysis

Table S1. Definitions of parameters used to illustrate household connectivity

| Network parameter | Definition |
| --- | --- |
| Degree | Measures the number of connections to and from each set of nodes (household, and water and grazing locations) in the six months preceding the study. In our network, connections between households and water and grazing locations were represented by directed edges. Put simply, households with more connections reflected higher levels of interaction at water and grazing locations within the network. |
| Betweenness | Betweenness centrality reflects the extent to which households within one village occupies strategic positions that influence connection to other villages in the network through the shortest paths. In our study, households with high weighted betweenness played a vital role in epidemiologically linking others in different villages and shaping the network's overall structure. Betweenness scores ranged from 0 (lowest centrality) to 1 (highest centrality). |
| Eigenvector centrality | Indirectly measures centrality based on how connected a node is to other nodes in the network. In our study, we used the eigenvector centrality to describe how connected a household is to other well-connected households - describing the importance of each household in the network A high eigenvector score means a household is connected to many others that are also highly connected. |

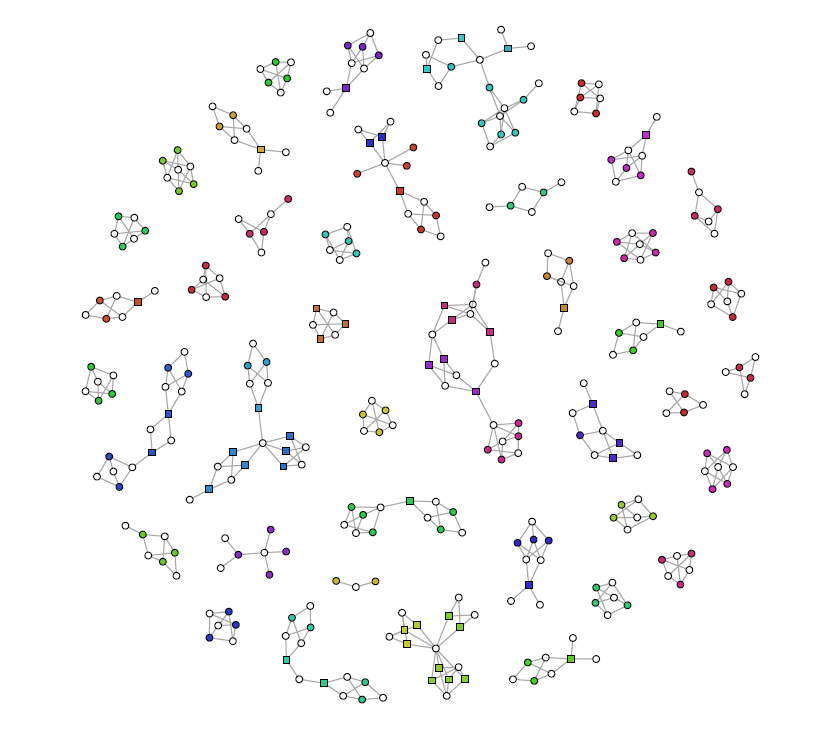

*Figure S2: A topological network depicting the overall connectivity among study households based on shared resource areas in The Gambia. The network is undirected and unweighted, with gray edges. Coloured circles denote individual households within specific villages, with varying hues representing different study villages. White circles indicate shared water and grazing locations used by households. Squares represent transhumant household herds, which seasonally migrate over long distances (up to 80 km), establishing connections with household herds across villages. The transhumant movements are directed towards the Gambia river valley from within the Sahelian eco-region.*

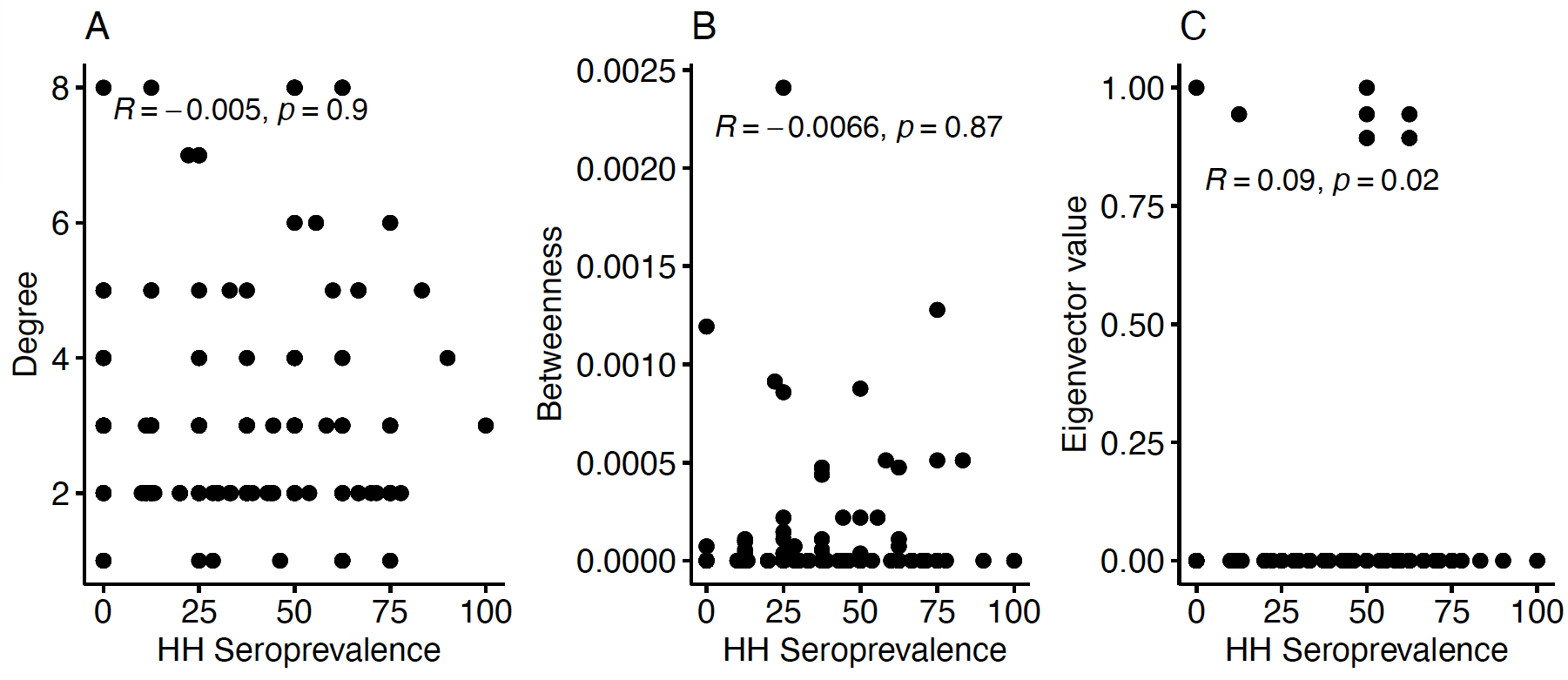
*Figure S3: Spearman's rank correlation analysis illustrating the* *relationship between household (HH) RVFV seropositivity in cattle and unweighted, normalized A: Degree, B: Betweenness, and C: Eigenvector value, derived from this study.*

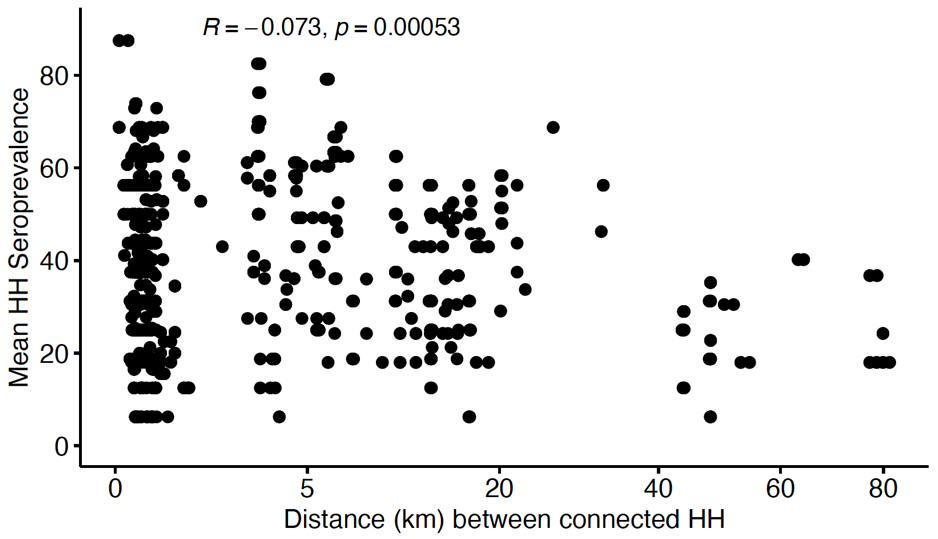

*Figure S4: Spearman's rank correlation analysis illustrating the relationship between the geographic distance separating connected households (HH) and their mean RVFV seropositivity of cattle in this study*

##### Epidemiological modelling

##### Estimation of model parameters

By assuming homogenous mixing of the cattle population and a dynamic process at endemic equilibrium, $R_{0,i}$ can be related to the proportion of the total cattle population in each eco-region that is susceptible, ${s^{*}}_{i}$, as follows:

$$R_{0,i}= \frac{1}{{s^{*}}_{i}} (1)$$

(Anderson and May, (1991). The proportion of susceptible cattle in each eco-region was approximated from the observed seroprevalence of each population, $F_{j}$, and the total number of cattle within the eco-region, $\tilde{N}_{i}$, as follows:

$${s^{*}}_{i} \sim1-\left( \frac{\sum_{k\epsilon\left\{ M,L,H \right\}} F_{k} N_{k}}{\tilde{N}_{i}} \right) (2)$$

However, $R_{0,i}$can also be approximated by:

$$R_{0,i}\sim\frac{\beta_{i} \tilde{N}_{i}}{\left( \mu+\gamma+\delta\right)} (3)$$

leaving $\beta_{i}$ as the remaining unknown.

The transmission rates during the wet and dry season were therefore estimated by:

$$\beta_{s,wet} \sim\psi\left( \frac{R_{0,s}\left( \mu+\gamma+\delta\right)}{N_{s,wet}} \right) (4)$$

$$\beta_{r,wet}\sim\frac{R_{0,r}\left( \mu+\gamma+\delta\right)}{N_{r,wet}} \left( 5 \right)$$

$$\beta_{r,dry} \sim\frac{R_{0,r}\left( \mu+\gamma+\delta\right)}{N_{r,dry}} (6)$$

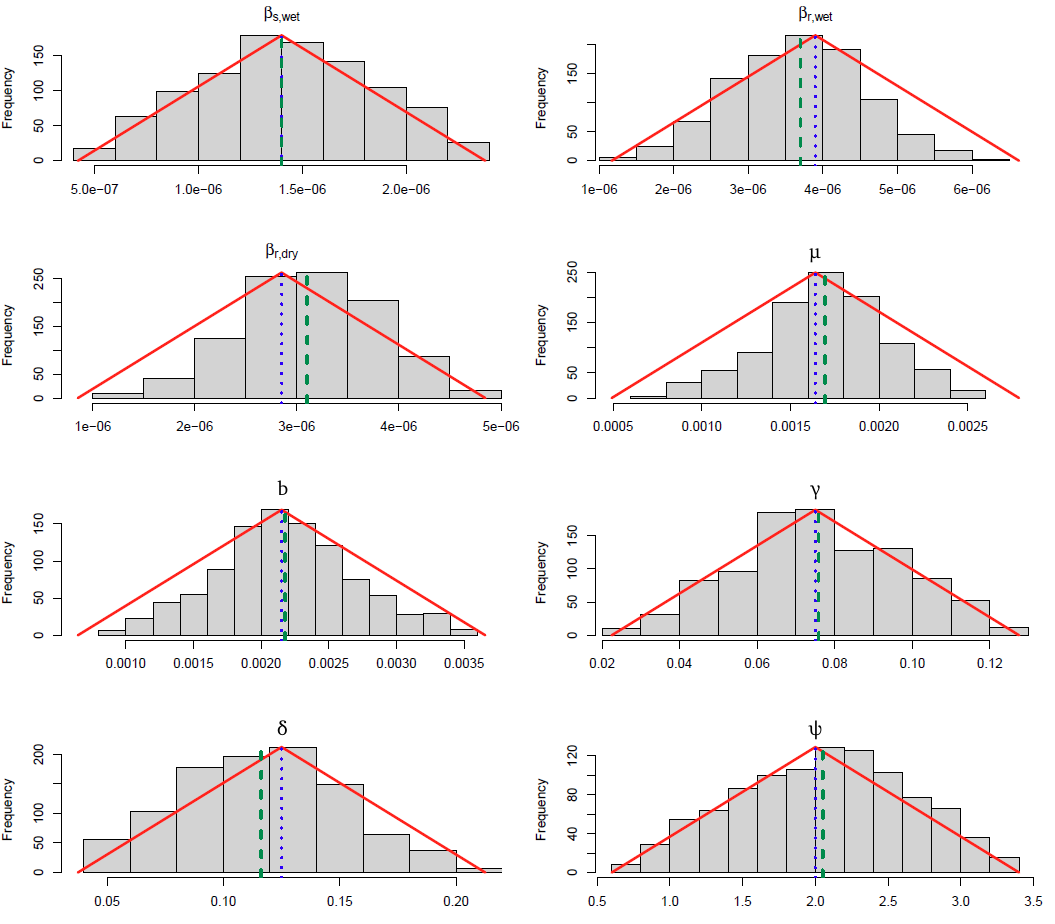
*Figure S5: Posterior distributions (gey bars) of the eight estimated parameters after fitting the SIR model to the observed seroprevalence using the ABC method.* $\beta_{s,wet}:$ *transmission parameter in the Sahelian eco-region in the wet season;* $\beta_{r,wet}$ *and* $\beta_{r,dry}$*: transmission parameter in the Gambia river region in the wet and dry seasons, respectively;* $\mu$*: natural death rate; b: per capita birth rate;* $\gamma$*: RVF-specific mortality rate;* $\delta$*: rate of recovery from RVF in cattle;* $\psi$*: scaling factor for* $\beta_{s,wet}$*. The red lines represent the triangular distribution of the parameter priors. Blue dotted lines: the preliminary parameter values used to generate the prior distribution. Green dashed lines: the mean values of the accepted posterior distribution from the ABC analysis.*

##### Ordinary differential equations

The ODEs that mathematically formalise the number of susceptible, infectious and recovered cattle of the T and M populations in the Sahelian eco-region during the wet season are as following:

$$\frac{dS_{T}}{dt}=bN_{T}-{\mu S}_{T}-\beta_{s,wet}S_{T}I_{T}-\beta_{s,wet}S_{T}I_{M} (7)$$

$$\frac{dI_{T}}{dt}=-{\left( \mu+\gamma+ \delta\right)I}_{T}+\beta_{s,wet}S_{T}I_{T}+\beta_{s,wet}S_{T}I_{M} (8)$$

$$\frac{dR_{H}}{dt}=\delta I_{T}-{\mu R}_{T} (9)$$

$$\frac{dS_{M}}{dt}=bN_{M}-{\mu S}_{M}-\beta_{s,wet}S_{M}I_{M}-\beta_{s,wet}S_{M}I_{T} (10)$$

$$\frac{dI_{M}}{dt}=-{\left( \mu+\gamma+ \delta\right)I}_{M}+\beta_{s,wet}S_{M}I_{M}+\beta_{s,wet}S_{M}I_{T} (11)$$

$$\frac{dR_{M}}{dt}=\delta I_{H}-{\mu R}_{M} (12)$$

The state of the L population in the Gambia river eco-region during the wet season after population and infection dynamics for cattle in the susceptible, infectious and recovered compartments was modelled by:

$$\frac{dS_{L}}{dt}=bN_{L}-{\mu S}_{L}-\beta_{r,wet}S_{L}I_{L} (13)$$

$$\frac{dI_{L}}{dt}=-{\left( \mu+\gamma+ \delta\right)I}_{L}+\beta_{r,wet}S_{L}I_{L} (14)$$

$$\frac{dR_{L}}{dt}=\delta I_{L}-{\mu R}_{L} (15)$$

The ODEs for the susceptible, infectious and recovered cattle of the T and L populations in the Gambia river valley during the dry season are as following:

$$\frac{dS_{T}}{dt}=bN_{T}-{\mu S}_{T}-\beta_{r,dry}S_{T}I_{T}-\beta_{r,dry}S_{T}I_{L} (16)$$

$$\frac{dI_{T}}{dt}=-{\left( \mu+\gamma+ \delta\right)I}_{T}+\beta_{r,dry}S_{T}I_{T}+\beta_{r,dry}S_{T}I_{L} (17)$$

$$\frac{dR_{T}}{dt}=\delta I_{T}-{\mu R}_{T} (18)$$

$$\frac{dS_{L}}{dt}=bN_{L}-{\mu S}_{L}-\beta_{r,dry}S_{L}I_{L}-\beta_{r,dry}S_{L}I_{T} (19)$$

$$\frac{dI_{L}}{dt}=-{\left( \mu+\gamma+ \delta\right)I}_{L}+\beta_{r,dry}S_{L}I_{L}+\beta_{r,dry}S_{L}I_{T} (20)$$

$$\frac{dR_{L}}{dt}=\delta I_{L}-{\mu R}_{L} (21)$$

The population dynamics in the M populations in the Sahelian eco-region during the dry season are modelled as following:

$$\frac{dS_{M}}{dt}=bN_{M}-{\mu S}_{M}-\beta_{s,dry}S_{M}I_{M} (22)$$

$$\frac{dI_{M}}{dt}=-{\left( \mu+\gamma+ \delta\right)I}_{M}+\beta_{s,dry}S_{M}I_{M} (23)$$

#####

$$\frac{dR_{M}}{dt}=\delta I_{M}-{\mu R}_{M} (24)$$

##### Estimation of region-specific RVFV force of infection

We determined the variation in $\lambda_{region,season}$ when the RVFV dynamics is at quasi-equilibrium as:

$$\lambda_{s,wet}=\beta_{s,wet}I_{s,M}(t)+\beta_{s,wet}I_{s,T}(t) (25)$$

$$\lambda_{r,wet}=\beta_{r,wet}I_{r,L}(t) (26)$$

$$\lambda_{r,dry}=\beta_{r,dry}I_{r,L}(t)+\beta_{r,dry}I_{r,T}(t) (27)$$

##### Estimation of the basic and seasonal reproduction numbers

The framework outlined below for estimating $R_{0,st}$ also applies for the estimation of $R_{0}$, with the only difference being the substitution of the number of susceptible cattle in each population at the end of each week, $S_{j}(t)$, with the fully susceptible cattle population at the start of the simulation, $N_{j}$.

$R_{0,st}$ was estimated as the spectral radius of the matrix **Q**:

$$\mathbf{Q= -F}\mathbf{T}^{\mathbf{-1}} (28)$$

$\boldsymbol{F}_{\boldsymbol{region,wet}}= \left[ \begin{matrix} \beta_{s,wet} S_{M}(t) & 0 & \beta_{s,wet} S_{M}(t) \\ 0 & \beta_{r,wet} S_{L}(t) & 0 \\ \beta_{s,wet} S_{T}(t) & 0 & \beta_{s,wet} S_{T}(t) \end{matrix} \right]$

$$\boldsymbol{F}_{\boldsymbol{region,dry}}= \left[ \begin{matrix} 0 & 0 & 0 \\ 0 & \beta_{r,dry} S_{L}(t) & \beta_{r,dry} S_{L}(t) \\ 0 & \beta_{r,dry} S_{T}(t) & \beta_{r,dry} S_{T}(t) \end{matrix} \right]$$

$$\boldsymbol{T}= \left[ \begin{matrix} -\left( \mu+\gamma+ \delta\right) & 0 & 0 \\ 0 & -\left( \mu+\gamma+ \delta\right) & 0 \\ 0 & 0 & -\left( \mu+\gamma+ \delta\right) \end{matrix} \right]$$

$$\boldsymbol{T}^{\boldsymbol{-1}}= \left[ \begin{matrix} -\frac{1}{\mu+\gamma+ \delta} & 0 & 0 \\ 0 & -\frac{1}{\mu+\gamma+ \delta} & 0 \\ 0 & 0 & -\frac{1}{\mu+\gamma+ \delta} \end{matrix} \right]$$

We then numerically solved for $R_{0,st}$ by calculating the leading eigenvalue of the matrix **Q**. $R_{0,st}$ values were calculated independently for each season.

##### Estimation of the per capita rate of decay of RVFV seropositivity

A version of our SIR model was formulated in terms of cattle age, $a,$ to predict RVFV age-seroprevalence profiles for each population subject to the estimated $\lambda_{region,season}$ for the Sahelian and Gambia river eco-regions. The differential equations which governed the seroprevalence dynamics in each population are as follows:

$$\frac{dS_{j}}{da}= - \lambda_{region, season}S_{j} (29)$$

$$\frac{dP_{j}}{da}= \left( \frac{\delta}{\delta+\gamma} \right)\lambda_{region,season}S_{j}- {\pi P}_{j} (30)$$

$$\frac{dD_{j}}{da}= \pi P_{j} (31)$$

Note: $\lambda_{region,season}$ signifies the force of infection in each eco-region during the wet or dry season.

##### Stochastic modelling and RVFV extinction risk

The local extinction risk in the transhumant (T) subpopulation is approximated as: $\frac{\sum V_{k}}{\sum Y_{k}}$

where:

$V_{k}$ = number of extinction events within the T subpopulation in the *k^th^* realisation.

$Y_{k}$ = duration (in years) of RVFV transmission in the *k^th^* realisation.

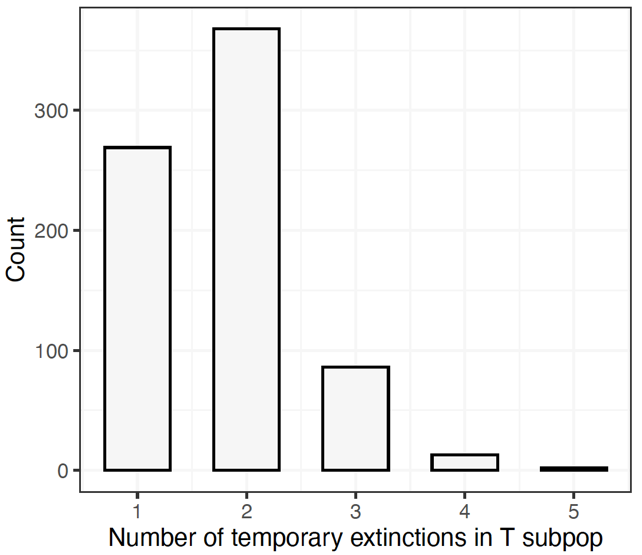

*Figure S6: The density plot showing the distribution of the number of local extinctions in the T subpopulation over an average transmission duration of 13.7 years. The x-axis represents the number of extinctions, and the y-axis represents the density (probability density function).*

##### Elasticity Analysis

Based on the comparison of the GLM and Loess fits, the relationship between the proportional change of the average predicted seroprevalence and proportional change of the posterior parameters was well approximated as linear.

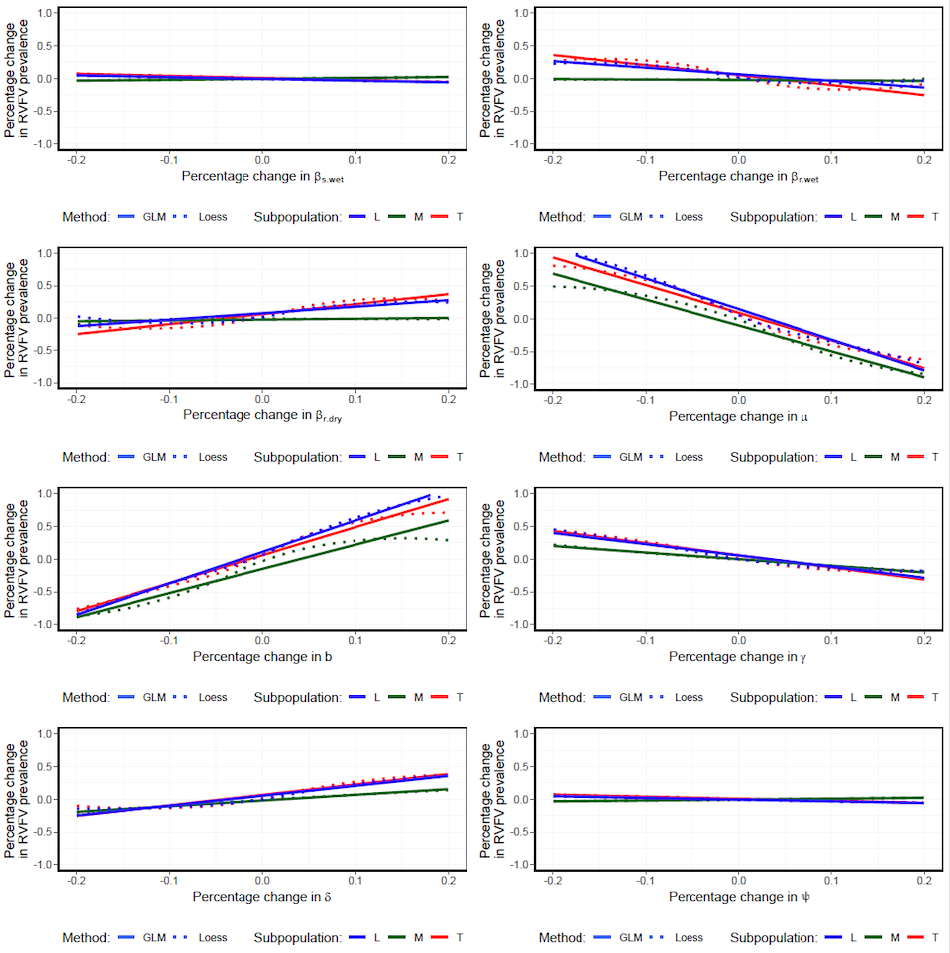
*Figure S7: The GLM (solid line) and Loess (dotted line) smoothing plots of the relationship between percentage change in predicted seroprevalence of each structured subpopulation (M, R and T) and percentage change in parameter values constrained within* $\pm$*20%* of the mean posterior value.
